# Cell-type specific transcriptomics reveals roles for root hairs and endodermal barriers in interaction with beneficial rhizobacterium

**DOI:** 10.1101/2022.05.09.491085

**Authors:** Eline H. Verbon, Louisa M. Liberman, Jiayu Zhou, Jie Yin, Corné M.J. Pieterse, Philip N. Benfey, Ioannis A. Stringlis, Ronnie de Jonge

## Abstract

Growth-promoting bacteria can boost crop productivity in a sustainable way. *Pseudomonas simiae* WCS417 is a well-studied bacterium that promotes growth of many plant species. Upon colonization, WCS417 affects root system architecture resulting in an expanded root system. Both immunity and root system architecture, are controlled by root-cell-type specific biological mechanisms, but it is unknown how WCS417 affects these mechanisms. Therefore, here, we transcriptionally profiled five *Arabidopsis thaliana* root cell types following WCS417 colonization. The cortex and endodermis displayed the most differentially expressed genes, even though they were not in direct contact with this epiphytic bacterium. Many of these genes are associated with reduced cell wall biogenesis, possibly facilitating the root architectural changes observed in WCS417-colonized roots. Comparison of the transcriptome profiles in the two epidermal cell types that were in direct contact with WCS417 – trichoblasts that form root hairs and atrichoblasts that don’t – imply functional specialization. Whereas basal expression levels of nutrient uptake-related genes and defense-related genes are highest in trichoblasts and atrichoblasts, respectively, upon exposure to WCS417 these roles revert. This suggests that root hairs participate in the activation of root immunity, further supported by attenuation of immunity in a root hairless mutant. Furthermore, we observed elevated expression of suberin biosynthesis genes and increased deposition of suberin in the endodermis in WCS417-colonized roots. Using an endodermal barrier mutant we show the importance of endodermal barrier integrity for optimal plant-beneficial bacterium association. Altogether, we highlight the strength of cell-type-specific transcriptional profiling to uncover “masked” biological mechanisms underlying successful plant-microbe associations.

## Introduction

Plants are sessile organisms that cannot move in response to environmental changes. Instead, they adapt to such changes by modifying the morphology and exudation of their roots or by activating a range of defense responses. The root system of the model plant *Arabidopsis thaliana* (Arabidopsis) consists of a primary root with branching lateral roots (Motte et al., 2019; Petricka et al., 2012). Modifications in the spatial configuration of roots, the root system architecture, are especially important for water and nutrient uptake (Koevoets et al., 2016; Li et al., 2016; Rogers and Benfey, 2015; Shahzad and Amtmann, 2017). Root structure is also vital for adaptation to different conditions. Plant roots are organized in concentric cycles consisting of different cell types, with the outer cell types (trichoblasts, atrichoblasts) being in contact with the environment and the inner ones (cortex, endodermis, pericycle, vasculature) being indispensable for nutrient/water transport between below- and aboveground plant tissues (Stassen et al., 2021; Wachsman et al., 2015).

Exudation of specialized plant metabolites and structural fortification of inner cell types such as the endodermis are essential for nutrient uptake from the soil and a balanced interaction with the microbial communities surrounding the roots, known as the microbiome (Kashyap et al., 2021; Pascale et al., 2020). Exudates such as coumarins can facilitate iron uptake from the soil but also shape the root microbiome (Harbort et al., 2020; Stringlis et al., 2018b). Fortification of the endodermis includes the coating of endodermal cells by a hydrophobic polymer, suberin, and the deposition of lignin-based structures to form the Casparian strip, in the junction between two adjacent endodermal cells (Barberon, 2017; Barberon et al., 2016; Geldner, 2013; Naseer et al., 2012). The amount of suberin deposition around endodermal cells is dynamically regulated during nutrient stresses and by the root microbiome (Barberon, 2017; Barberon et al., 2016; Salas-Gonzalez et al., 2021). An extra level of plant adaptation is achieved via the modification of root system architecture in response to beneficial soil micro- organisms (Vacheron et al., 2013; Verbon and Liberman, 2016). In Arabidopsis, the number and/or length of lateral roots and root hairs increase in response to different rhizobacteria and fungi (Contreras-Cornejo et al., 2009; Lopez-Bucio et al., 2007; Vacheron et al., 2018; Zamioudis et al., 2013). All the above mentioned chemical, morphological or structural modifications of roots towards the root microbiome rely on the prompt perception of microbes or their defense-eliciting molecules (Microbe-Associated Molecular Patterns or MAMPs). These changes ultimately allow plants to maintain a beneficial interaction with their microbiome and avoid colonization by unwanted and potentially harmful microbes (Beck et al., 2014; Colaianni et al., 2021; Hacquard et al., 2017; Millet et al., 2010; Stringlis et al., 2018a; Teixeira et al., 2019; Wyrsch et al., 2015).

Studies on the interaction between Arabidopsis and the beneficial rhizobacterium *Pseudomonas simiae* WCS417 (WCS417) unearthed different aspects of the interplay between plants and their associated beneficial microbes (Pieterse et al., 2021). WCS417 stimulates Arabidopsis growth (Berendsen et al., 2015; Zamioudis et al., 2013) and induces systemic resistance against many pathogens in Arabidopsis and several crop species (Pieterse et al., 1996; Pieterse et al., 2014). Arabidopsis responds to root colonization by WCS417 by inhibiting primary root growth and increasing the number of lateral roots and root hairs (Stringlis et al., 2018a; Zamioudis et al., 2013). The increased number of lateral roots upon WCS417 colonization is due to an increase in lateral root initiation events, observed as an increased number of lateral root primordia, and increased outgrowth of these primordia (Zamioudis et al., 2013). Lateral roots originate from pericycle cells, a cell layer surrounding the vasculature, and subsequently force their way through the endodermis, cortex, and finally the epidermis, to protrude from the primary root (Du and Scheres, 2018; Malamy and Benfey, 1997; Moller et al., 2017; Otvos and Benkova, 2017). The plant hormone auxin is important for all phases of lateral root development (Du and Scheres, 2018). In line with this, the increase in lateral root number in response to WCS417 is dependent on auxin signaling (Zamioudis et al., 2013). Similarly, the WCS417-mediated increase in root hair number is dependent on auxin signaling (Zamioudis et al., 2013). In Arabidopsis, root hairs are formed by specialized cells in the epidermis: the trichoblasts. Together with the other cell type in the epidermis, the atrichoblasts, they form the outermost root cell layer (Gilroy and Jones, 2000; Ryan et al., 2001; Vissenberg et al., 2020). The activity of several transcription factors, including TRANSPARENT TESTA GLABRA (TTG), CAPRICE (CPC) and WEREWOLF (WER), and the spatial localization of the cells, with cells located over two cortical cells becoming trichoblasts, regulate whether trichoblasts or atrichoblasts are formed (Vissenberg et al., 2020). In response to WCS417, the increased number of root hairs is due to an increased number of cortical cells and therefore an increased number of cells becoming trichoblasts (Zamioudis et al., 2013). A root system with a greater number of lateral roots and/or root hairs can mine more soil for nutrients, has a larger surface to facilitate colonization by plant growth-promoting rhizobacteria (PGPR) (Lugtenberg and Kamilova, 2009; Vacheron et al., 2013), and has greater potential to release nutrient-mobilizing exudates (e.g. Fe-chelating coumarins) (Robe et al., 2021).

Establishment and maintenance of beneficial plant-microbe interactions requires a fine MAMP-triggered immunity (MTI) which, when left unchecked, inhibits growth (Ma et al., 2021; Teixeira et al., 2021). Previous studies demonstrated that WCS417 can repress part of the root defense responses (Millet et al., 2010; Stringlis et al., 2018a), probably via the production of gluconic acid (Yu et al., 2019b). In recent years, many studies have demonstrated that the different cell types of the root can mount defense responses of varying levels depending on the MAMP and the responsible microbe colonizing the roots (Rich-Griffin et al., 2020a; Salas-Gonzalez et al., 2021; Wyrsch et al., 2015; Zhou et al., 2020). These studies suggest that by compartmentalizing detection of microbes and activation of defense responses, the plant can maintain a proper growth - defense balance, avoiding costly and/or late defense activation (Teixeira et al., 2019; Yu et al., 2019a).

The structure of the Arabidopsis root system is defined by the distinct biological functions of each of its cell types. In parallel, specialized responses activated in each cell type upon microbial colonization allow plants to grow optimally in microbe-rich environments. Our previous studies on whole roots provided us with global information on the interaction between WCS417 and Arabidopsis (Stringlis et al., 2018a; Verhagen et al., 2004; Zamioudis et al., 2014). However, cell-type-specific transcriptomics can reveal which cell types respond to WCS417 most strongly or quickly, what responses are activated in each cell type, and how these responses contribute to successful colonization and subsequent effects on root architecture and the establishment of a mutualistic interaction (Rich-Griffin et al., 2020b). We used a set of fluorescent marker lines to isolate trichoblast, atrichoblast, cortical, endodermal and vasculature cells with fluorescence-activated cell sorting (FACS) (Birnbaum et al., 2005; Birnbaum et al., 2003; Brady et al., 2007a). To build a map of gene expression changes in the root, we transcriptionally profiled these cell populations after colonization by WCS417. Our data show distinct cell-type specific responses to WCS417 exposure. The most dramatic changes are seen in the cortex and endodermis where genes involved in cell wall reorganization reflect the morphological observations of increased lateral root formation. Additionally, endodermal cells increase their protective barrier in response to WCS417 by increasing suberin biosynthesis. We also found evidence for functional specialization of the root epidermal cell types indicating a prominent role for trichoblasts in nutrient uptake under control conditions and activation of immunity upon bacterial treatment. We suggest that root hairs act as antennae for microbial signals and the generation of downstream responses.

## Results

### WCS417 rapidly induces root developmental changes

PGPR can affect plant root system architecture (Vacheron et al., 2013; Verbon and Liberman, 2016). In accordance with previous reports (Stringlis et al., 2018a; Zamioudis et al., 2013), WCS417 inhibits primary root length and increases the total number of lateral roots after seven days of co-inoculation in a dose-dependent manner (Figure S1). Dose-dependency of PGPR-mediated increases in plant growth and resistance to disease is a common phenomenon (Asari et al., 2017; Farag et al., 2013; Raaijmakers et al., 1995; Ryu et al., 2003). After only two days of co-inoculation we observed increased formation of lateral roots when 10^7^ or more bacteria were applied per row of plants (Figure S1). Therefore, the effects of WCS417 on root growth are visible at 48 h and these effects are dose-dependent. We reasoned that by studying this timepoint using cell-type-specific transcriptomics, we would capture the transcriptional events in different cell types underlying early plant modifications in response to WCS417 and identify processes involved in the establishment of a beneficial plant-microbe interaction.

### Cell-type-specific transcriptional profiling of the Arabidopsis root

To create a spatial map of root transcriptional changes in response to colonization by WCS417, we isolated several root cell types using FACS. First, we confirmed that WCS417 does not affect the expression pattern of *GREEN FLUORESCENT PROTEIN* (*GFP*) when driven by the cell-type-specific promotors *WEREWOLF* (*WER*; atrichoblast), *COBRA-LIKE 9 (COBL9*; trichoblast), *315* (cortex), *SCARECROW* (*SCR*; endodermis), or truncated *WOODENLEG* (*WOL*; vasculature) (Figure 1A). Subsequently, we grew the transgenic lines carrying these promotor-GFP fusions under high-density conditions and exposed them to WCS417. Two days after inoculation with WCS417, we harvested the roots, performed FACS and isolated RNA (Figure 1B).

**Figure 1.**
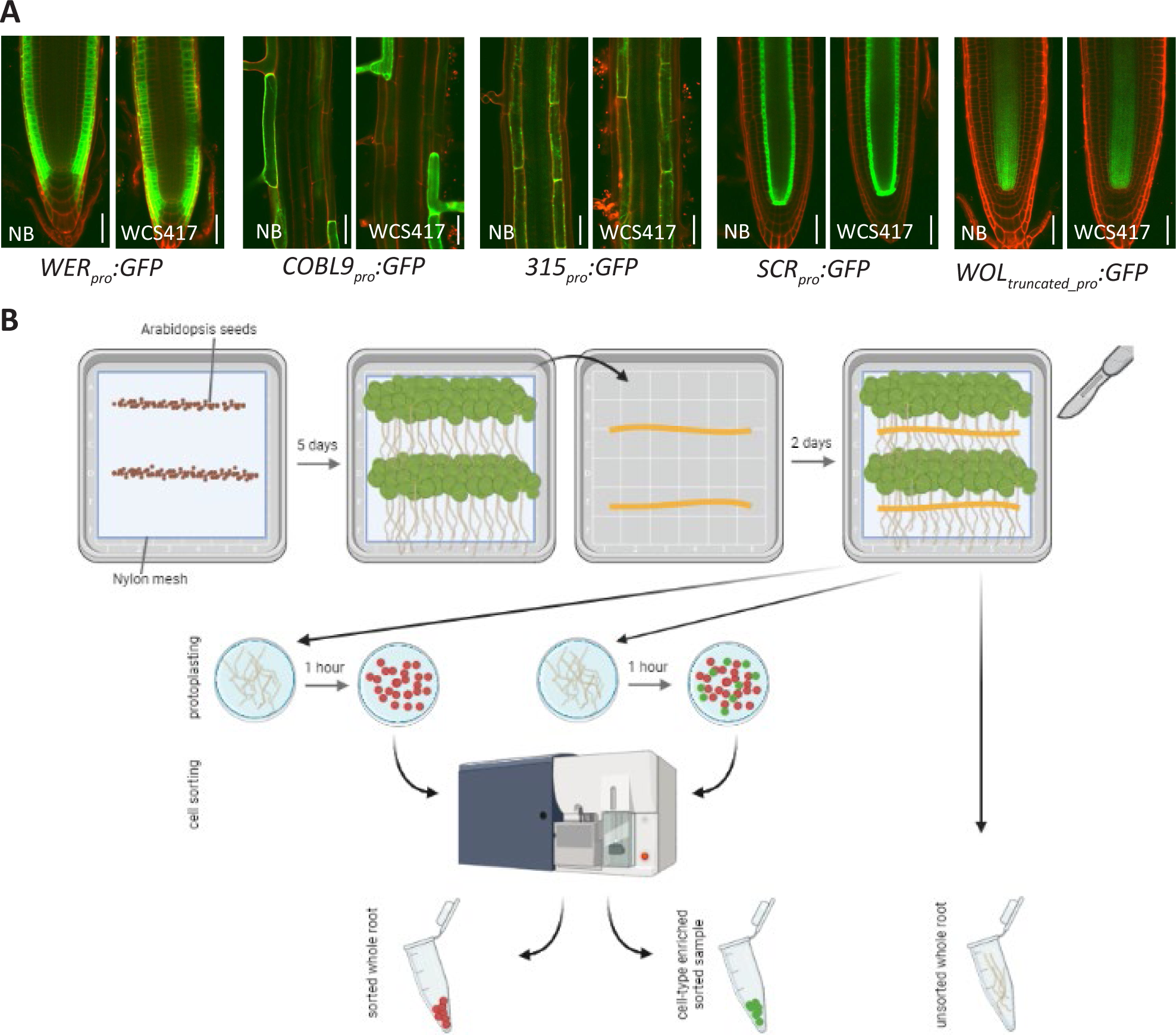
Exposure of five transgenic plant lines to WCS417 to obtain cell-type-specific samples by performing fluorescence-activated cell sorting (FACS). A) Sterile and WCS417-exposed plants from the transgenic plant lines *WEREWOLF_pro_:GFP* (*WER*: immature epidermis and atrichoblast), *COBRA-LIKE9_pro_:GFP* (*COBL9*: trichoblast), *315_pro_:GFP* (*315*: cortex), *SCARECROW_pro_:GFP* (*SCR*: endodermis), and *WOODENLEG_truncated_pro_:GFP* (*WOL*: vasculature). Pictures of the seedlings were taken all along the root from day five (day of bacterial inoculation) till day seven. GFP settings were kept the same between bacteria-exposed and sterile-grown plants. Representative images are shown. **B**) Experimental design used to obtain WCS417-treated and control samples enriched for one out of five root cell types. Sterilized and vernalized Arabidopsis seeds were sown on 1 × MS 1% sucrose plates and left to grow in long-day conditions. Five days later, half of the plants of each line were transferred on their mesh onto 1 × MS 1% sucrose plates with WCS417. Plants were left to grow for a further two days of growth before root harvest. Wild-type Col-0 roots were either directly flash- frozen (unsorted control) or protoplasted and put through the cell sorter, collecting non-fluorescent cells (sorted control). Transgenic lines with cell-type specific *GFP* expression were similarly protoplasted and put through the cell sorter for fluorescence-activated cell sorting (FACS).

To determine the success of the sorting procedure, we checked the expression of the marker genes *WER*, *COBL9*, *315*, *SCR* and *WOL* in our transcriptomic dataset (Figure 2A). The expression of each of these markers should be highest in the FACS samples obtained from the transgenic plant lines in which the corresponding promotor was used to drive *GFP* expression. Indeed, the expression of *WER*, *COBL9*, *315* and *SCR* is highest in the samples obtained from their respective lines (Figure 2B). The expression of well-established vasculature marker genes, namely *INCURVATA4, SHORTROOT* and *ZWILLE* is enriched as expected in the *pWOL_truncated_pro_:GFP* line, but the expression of *WOL* is not enriched (Figure 2C). This suggests that only the truncated *WOL* promotor, and not the full promotor, is cell-type-specific.

**Figure 2.**
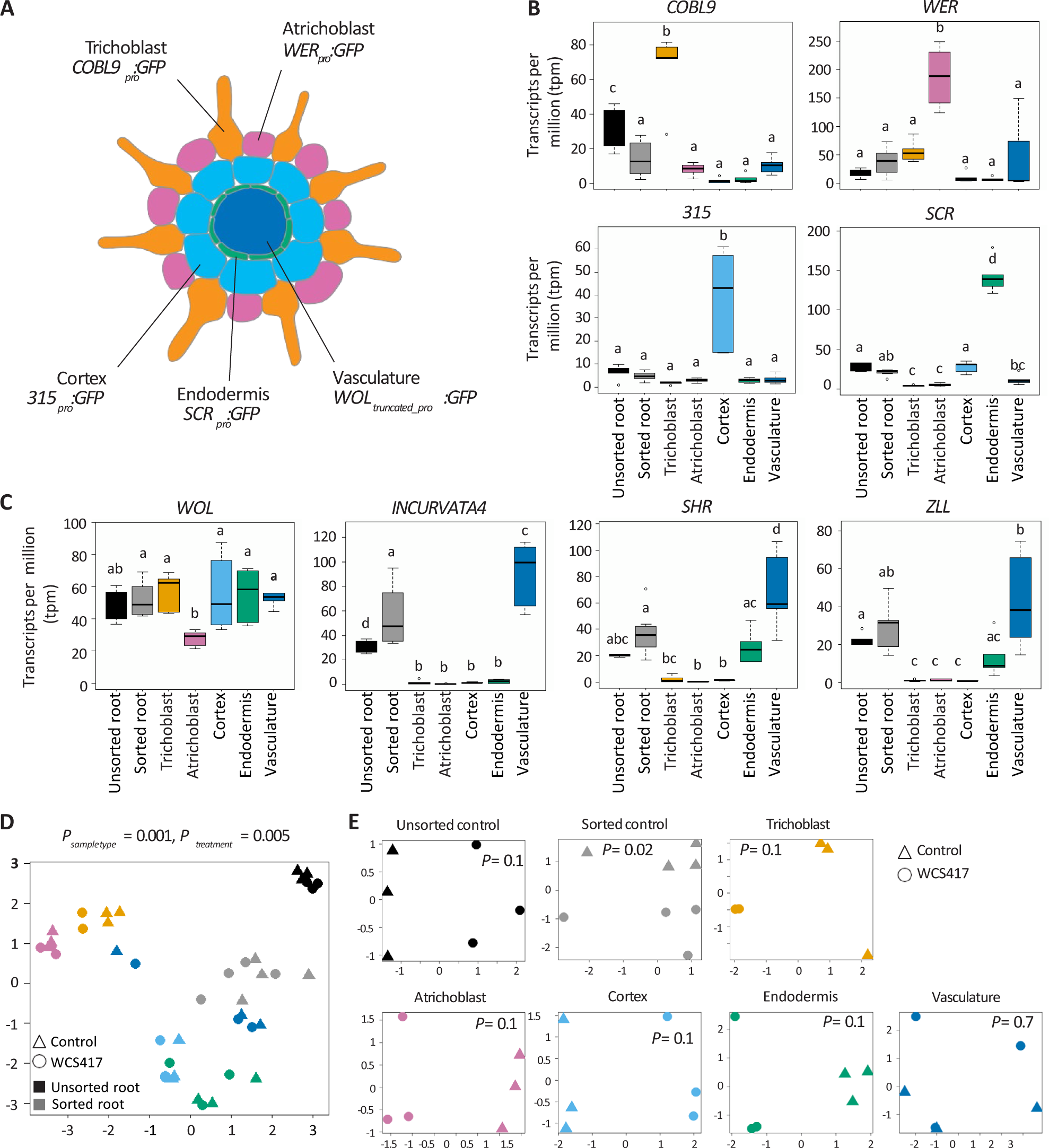
Gene expression differences among the samples reflect Arabidopsis root development patterns. **A**) Schematiccross section of the Arabidopsis root, with each cell type labeled with the *promoter::GFP* fusion that was used to enrich samples for that cell type by FACS. **B**) mRNA levels of the marker genes *COBL9*, *WER*, *315*, and *SCR*. **C**) mRNA levels of the marker gene *WOL,* and of the vasculature-specific genes *INCURVATA4, SHR*, and *ZLL*. Data was analyzed with an ANOVA test followed by the Tukey post hoc test in R (p-value <0.05). GFP: green fluorescent protein, COBL9: COBRA-LIKE 9, WER: WEREWOLF, SCR: SCARECROW, WOL: WOODENLEG, SHR: SHORTROOT, ZLL: ZWILLE. Multidimensional scaling (MDS) plot of counts (log scale) per million of all samples (**D**) and per cell type (**E**). WCS417-exposed samples are represented by circles, control, untreated samples by triangles. Colors in panel B-E correspond to the color scheme of the schematic in panel A. Black samples represent the unsorted wild-type roots, grey represents the sorted wild-type roots. InpanelE,‘*P*’represents the p-value of the WCS417-treatment effect.

To study the global similarities and dissimilarities among samples and treatments, we performed multidimensional scaling on gene expression levels. The transcriptional profiles cluster by sample type (*P_sample type_* = 0.001; Figure 2D). The cortical and endodermal cells cluster close together, as do the two epidermal cell types. This is in line with the known development of the Arabidopsis root, in which the cortex and endodermis develop from a shared stem cell population, as do the trichoblasts and atrichoblasts (Dolan et al., 1993; Van den Berg et al., 1995). In addition to the sample-type effect, we find an effect of bacterial treatment on gene expression (*P_treatment_* = 0.005; Figure 2D). When comparing gene expression patterns of samples within sample types, each cell type except the vasculature clusters primarily based on bacterial treatment (Figure 2E).

Next, we determined which genes are differentially expressed (DEGs) in response to WCS417 in each of the cell types compared to untreated roots (Table S2). The number of DEGs differs greatly among the cell types, ranging from 30 in the vasculature to 1,109 in the cortex (Figure 3A). Interestingly, the cortical and endodermal cells, which do not interact directly with the strictly epiphytic WCS417 bacterium, displayed the largest number of DEGs (1,109 and 815, respectively), while the trichoblast and atrichoblast cells, which are in direct contact with WCS417 displayed much less DEGs (469 and 137, respectively). Apart from a quantitative difference, the response is also qualitatively different between cell types: of the 1,862 DEGs across all five cell types, 72% are affected in only one cell type and only six genes are affected in all cell types (Figure 3B). Notably, the majority of genes affected in only a single cell type are not identified as differentially expressed in the whole root, while most genes affected in four or five cell types are identified as either up- or down-regulated in whole roots (Table S3). In contrast, the majority of the genes found to be up- or down-regulated in the sorted or unsorted control were identified as differentially expressed in response to WCS417 in one or more cell types (Table S4). In conclusion, genes affected in only single cell types are often not identified as differentially expressed in the whole root. This explains the higher number of identified DEGs in the cell-type-specific data set as compared to the sorted whole root (1,862 genes versus 270 genes; Figure 3A).

**Figure 3.**
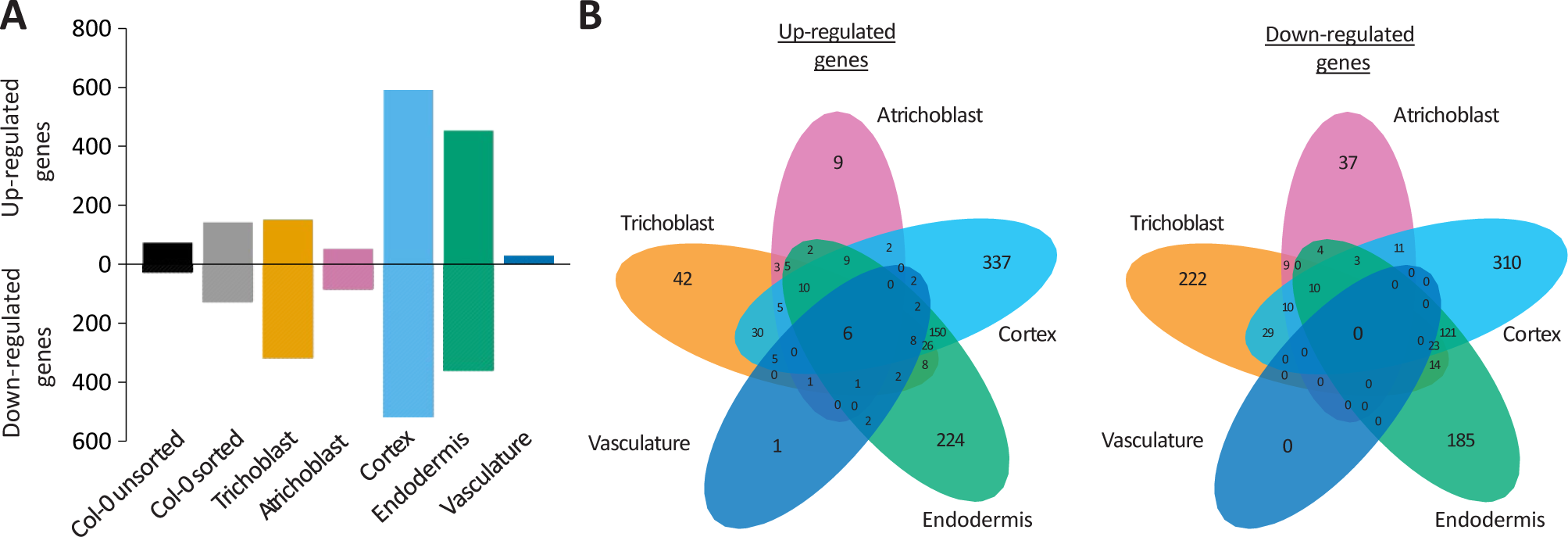
Root cell types have unique responses to root colonization by WCS417. **A**) Number of differentially expressed genes upon WCS417 application found in the respective samples (false discovery rate (FDR) < 0.1; -2 < log2FC > 2). **B**) Venn diagrams showing the overlap in genes affected by WCS417 treatment in the five studied cell types.

### Distinct specializations of the trichoblasts and atrichoblasts

To identify the biological processes affected by WCS417 colonization in the different cell types, we conducted biological process gene ontology (GO) term enrichment analyses on the DEGs (Table S5-S14). Most significant among the up-regulated DEGs in the trichoblasts, cortex, endodermis and vasculature are processes related to defense and immunity (Table 1).

Interestingly, atrichoblasts do not respond to WCS417 with defense activation, but with activation of ion transport (Table 1). This suggests that the two cell types directly in contact with WCS417 activate distinct biological processes, as could be expected from the limited overlap in DEGs between these cell types (Figure 3B and 4A). To further analyze these differences, we examined the expression of all genes within the GO terms defense response (GO:0006952) and ion transport (GO:0009267) that are differentially expressed in one or both epidermal cell types. Based on their expression levels, the genes involved in the defense response form five clusters (Table S15, Figure 4B, left). The two largest gene clusters (cluster 2 and 4) consist of genes that are induced by WCS417. Interestingly, the expression of these genes is observed in both WSC417-treated epidermal cell types. In contrast, the expression levels in control conditions are distinct, with lower gene expression in trichoblasts (Figure 4B, Table S17). When analyzing the expression patterns of ion transport-related genes, clusters 1, 3 and 5 contain genes that are up-regulated in response to WCS417 in both cell types (Figure 4B, Table S16). The differences between trichoblasts and atrichoblasts are clear in clusters 2, 4 and 6, which contain genes that are only expressed in trichoblasts in control conditions (Figure 4B, right).

**Figure 4.**
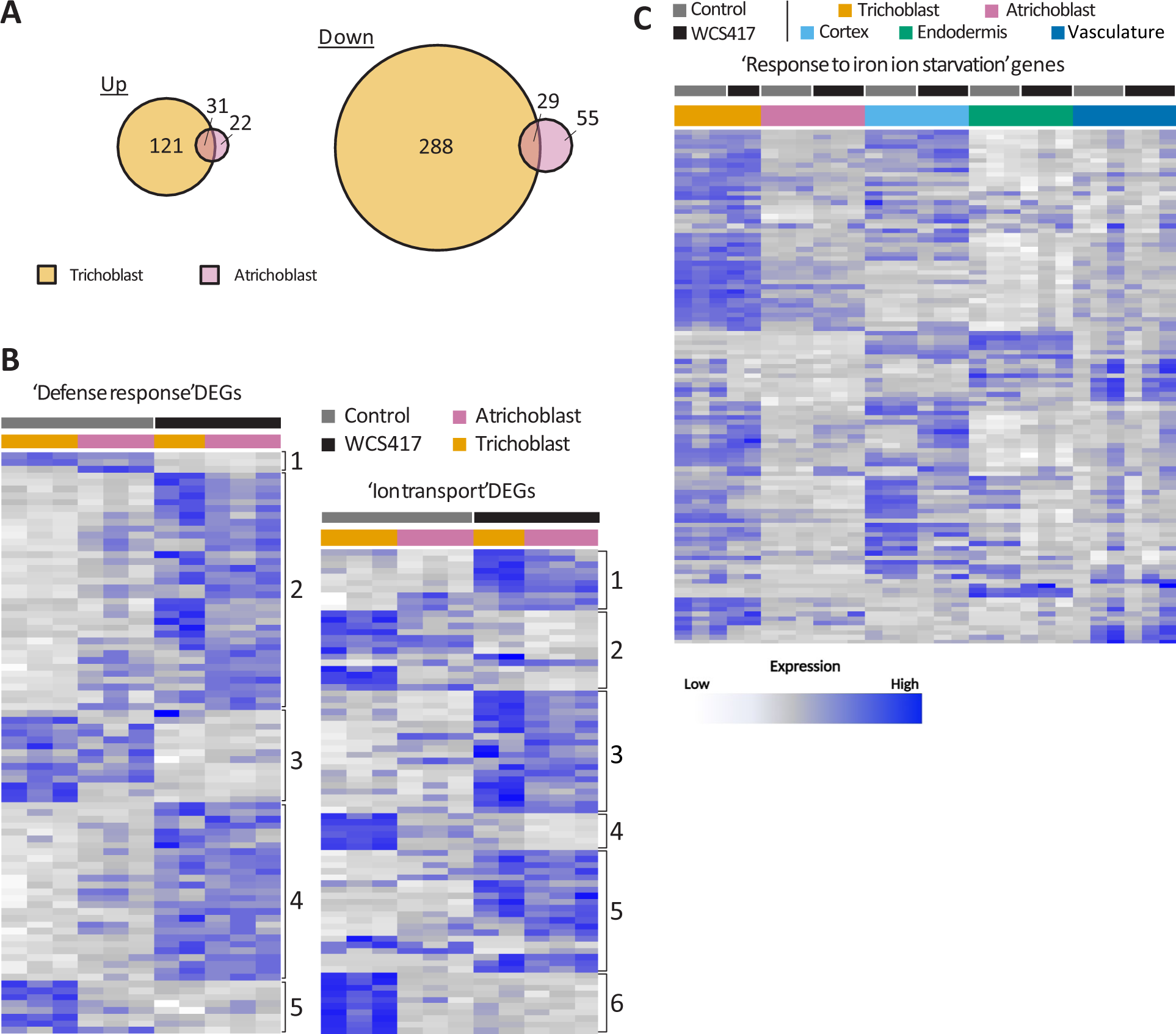
Specialization of the trichoblast in nutrient uptake and the atrichoblast in defense. **A**) Overlap of the DEGs in response to WCS417 in the trichoblasts and atrichoblasts. **B**) Heatmap of the expression of genes associated with the GO term defense response (GO:0006952, left) and the GO term ion transport (GO:0006811, right) that are differentially expressed in either or both the trichoblast and atrichoblast. Cluster numbers are based on visual differences in gene expression patterns. **C**) Heatmap of the expression of all genes associated with the GO term response to iron ion starvation (GO:0010106). Heatmaps are scaled by row. Gene expression is shown as the normalized log-counts-per-million, with low gene expression in white, and high expression in dark blue.

**Table 1.**
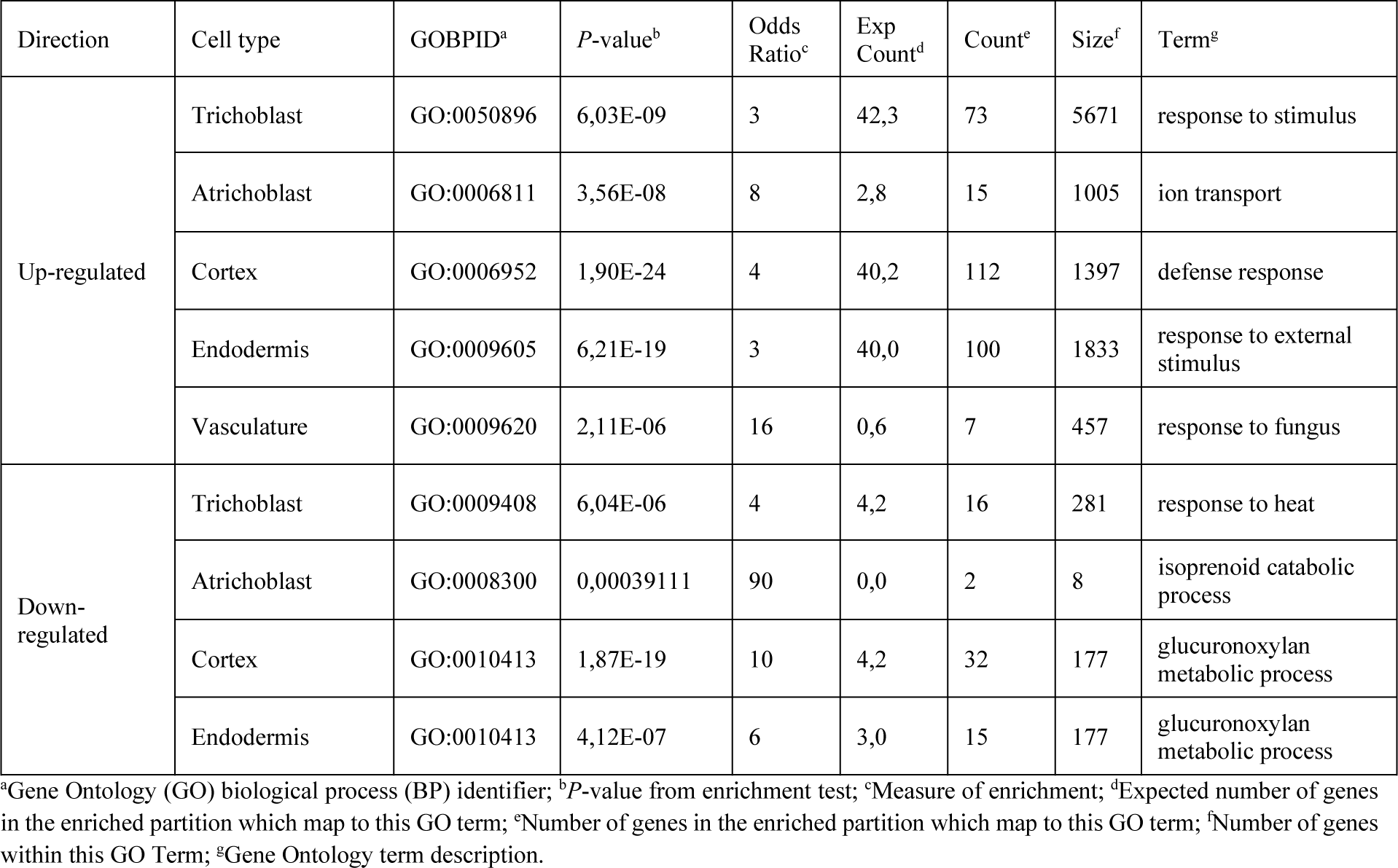
Top GO terms enriched in the WCS417-affected gene lists based on *p*-value.

These results suggest that distinct responses of trichoblasts and atrichoblasts to WCS417 are, in part, due to differences in basal gene expression patterns. To test this hypothesis, we investigated DEGs between the control, untreated, trichoblasts and atrichoblasts (Table S17), and identified enriched GO terms. Genes that are expressed higher in the atrichoblasts in control conditions are enriched for genes involved in RNA modification or processing, defense responses, response to hypoxia, response to salicylic acid and glucosinolate biosynthesis (Table S18). Genes that are expressed higher in trichoblasts are enriched for genes associated with the response to ion starvation, root hair differentiation, cell maturation, cell wall biosynthesis, root hair elongation, coumarin biosynthesis, (cell) growth, and the response to brassinosteroids / auxin / cytokinin (Table S19). Among these latter processes, iron ion starvation (GO:0010106) is enriched the most. Analysis of the expression of the genes involved in this process in all cell types shows that the expression of the majority of the genes involved in the response to iron ion starvation is primarily found in trichoblasts and, to a lesser extent, the cortex (Figure 4C, Table S20). This supports previous studies showing cortex- and epidermis-specific expression of the genes *BGLU42* and *IRT1*, both known to be involved in the iron deficiency response (Vert et al., 2002; Zamioudis et al., 2014). Recent data further support the importance of trichoblasts and root hairs for nutrient uptake and for the accumulation of iron-mobilizing coumarins (Robe et al., 2021; Tanaka et al., 2014). Thus, defense gene activity and expression levels of genes involved in nutrient uptake and root hair elongation are major differentiating factors between trichoblasts and atrichoblasts in control conditions in our experiment.

### Root hairs act as “antennae” for the perception of microbial signals and affect plant responses to WCS417

We found that while basal expression of defense-related genes, i.e., the expression in control (untreated) conditions, was the highest in atrichoblasts, upon exposure to WCS417, particularly trichoblasts appear to activate defense, illustrated by the enrichment for defense in these cells, specifically (Figure 4B, Table 1). This could indicate that cell types destined for the formation of root hairs (trichoblasts) might be more sensitive to microbial signals and this sensitivity could affect plant responses to microbes. To test this hypothesis, we assessed the effect of WCS417 on two mutants with contrasting patterns of root hair formation, *cpc* that cannot form root hairs and *ttg1* where most cells of the epidermis produce root hairs (Vissenberg et al., 2020). We grew Arabidopsis wild-type Col-0 plants and the *cpc* and *ttg1* mutants in plates containing 10^5^ colony-forming units (CFU) · ml^-1^ WCS417 based on a protocol developed by Paredes et al. (2018) and measured growth-promotion traits and levels of root colonization. Interestingly, the beneficial effects of WCS417 were less pronounced on both mutants as compared to wild-type plants, since relative changes in fresh shoot weight, primary root length and number of lateral roots were significantly lower (Figure 5A, B, C and Figure S2). Nevertheless, WCS417 colonization was comparable between wild-type and mutant roots (Figure 5D). We then reasoned that the number of root hairs might also affect defense responses to WCS417 and the bacterial MAMP flg22. For this, we tested the expression of WCS417 and/or flg22-responsive marker genes *MYB51, CYP71A12*, *PRX33* and *LECRK-IX.2* (Millet et al., 2010; Stringlis et al., 2018a). MYB51 and CYP71A12 have roles in indole glucosinolate and camalexin biosynthesis respectively (Millet et al., 2010), PRX33 is a cell wall peroxidase involved in the generation of reactive oxygen species (ROS) during defense activation (Kaman-Toth et al., 2019) and LECRK-IX.2 is a positive regulator of MTI (Luo et al., 2017). At 6 h after treatment, flg22 led to significant upregulation of all tested genes in roots of wild-type plants (Figure 5E-H), while WCS417 induced only the expression of *PRX33* (Figure 5H). Interestingly, flg22 didn’t affect the expression of any of the genes in the root hairless mutant *cpc*, and WCS417 caused only a slight induction of *PRX33* (Figure 5E-H). On the other hand, in roots of *ttg1* that produces more root hairs than the wild-type (Figure S2), *CYP71A12* and *PRX33* were upregulated to levels comparable to wild-type roots following flg22 treatment (Figure 5F, H). Strikingly, in *ttg1* the expression of *MYB51* and *LECRK-IX.2* was considerably higher in response to WCS417 and flg22 as compared to wild-type and *cpc*, but also under basal conditions (Figure 5E, G). Overall, it appears that the presence of root hairs affects the expression of MTI-related genes and the beneficial effects by WCS417 in mutants with altered root hair density are less pronounced.

**Figure 5.**
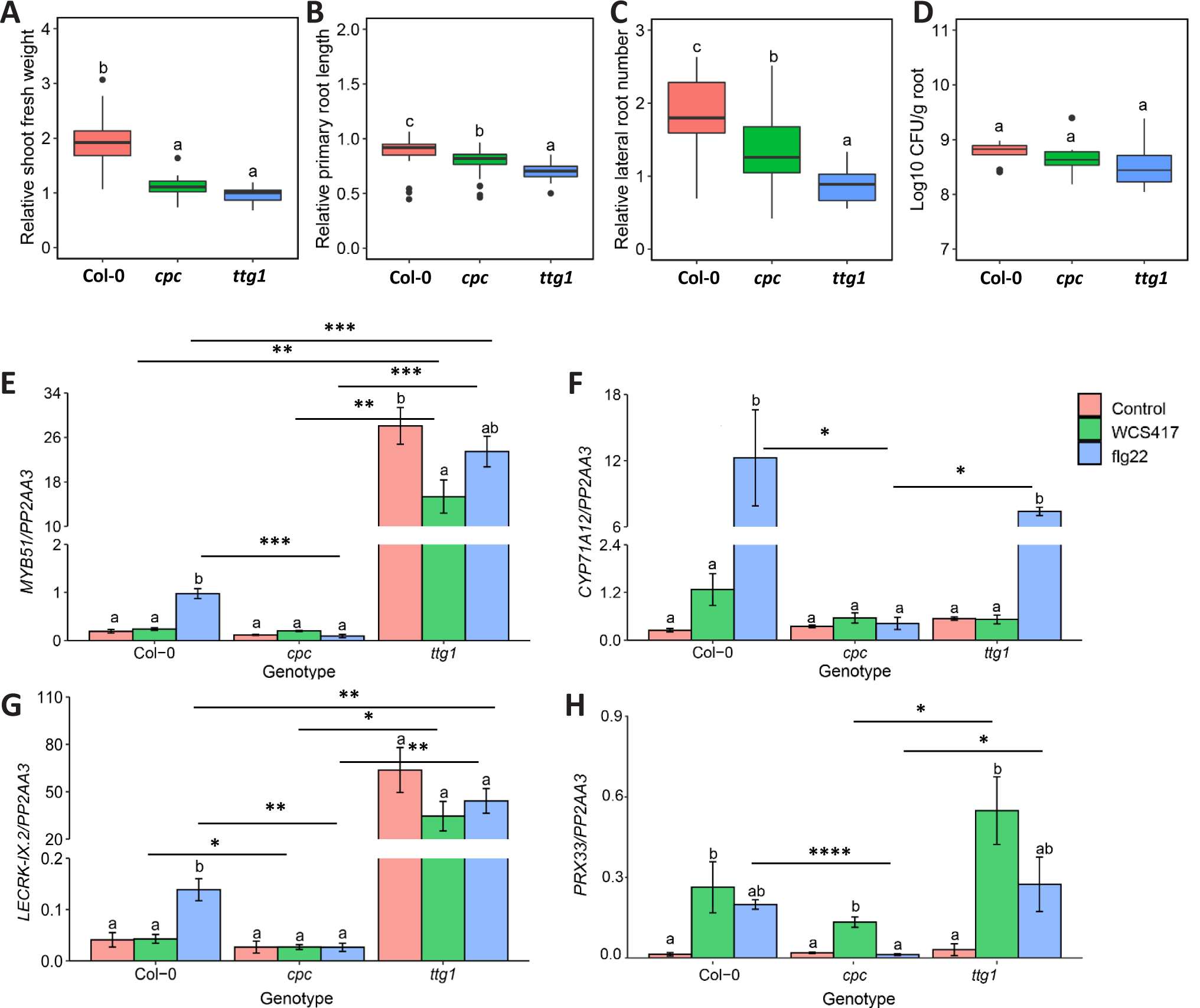
Arabidopsis root hair mutants display differential responses to WCS417. Relative **A**) shoot fresh weight, **B**) primary root length, **C**) lateral root number, and **D**) colonization levels of WCS417 on roots of Col-0, *cpc* and *ttg1* genotypes at 7 days after seedlings were transferred to plates with 5 Hoagland medium (0% sucrose) containing 10 CFU · ml^-1^ WCS417. Different letters indicate statistically significant differences across genotypes (One-way ANOVA, Tukey’s test; *P* < 0.05). In the case of growth parameters n= 30 and in case of root colonization n= 6. (**E – H**) Expression levels of *MYB51*, *CYP71A12*, *LECRK-IX.2* and *PRX33* as quantified by qRT-PCR. Expression was tested in roots of 8-day-old seedlings at 6 h after inoculation with WCS417 (OD_600_equal to 0.1, 10 CFU · ml^-1^) or treated with 1 μM flg22. Error bars represent SEM. Different letters represent statistically significant differences among control, WCS417 and flg22 in same plant genotype (One-way ANOVA, Tukey’s test; *P* < 0.05, n= 3-4). Asterisks indicate significant differences across genotypes that received the same treatment (WCS417 or flg22) (Student’s *t*-test; * *P* < 0.05, \*\**P* < 0.01, \*\*\**P* < 0.001,\*\*\*\**P* < 0.0001).

### WCS417 might facilitate lateral root formation by loosening cell walls of cell layers overlaying lateral root primordia

The most significant biological process among the down-regulated DEGs is the glucuronoxylan metabolic process (GO:0010413) in the cortex and endodermis (Table 1). Glucuronoxylan metabolic process is important for cell wall biosynthesis. In addition to this GO term, many other GO terms related to cell wall biosynthesis are enriched in down-regulated DEGs and, conversely, cell wall disassembly (GO:0044277) is enriched among the up-regulated DEGs in these cell types (Table S9-S12). Cell wall remodeling and cell volume loss in the cortex and endodermis are known to be required to accommodate emerging lateral roots and are possibly even required for the initiation of lateral root primordia (Stoeckle et al., 2018; Vermeer et al., 2014). Among the genes upregulated in the cortex and endodermis that are involved in cell wall disassembly are genes which encode polygalacturonases that have been shown to be expressed at the site of lateral root emergence. They are implicated in cell separation, possibly to accommodate emerging lateral roots (Ogawa et al., 2009). Up-regulation of these and other genes involved in cell wall disassembly and down-regulation of genes involved in cell wall biosynthesis in the cortex and endodermis might therefore be an integral part of the molecular and physiological changes that take place in response to WCS417, which lead to the observed increase in the number of lateral roots (Figure S1B and (Zamioudis et al., 2013)).

### WCS417 induces suberin biosynthesis in endodermal cells

Defense in general is the most significant process affected in our cell-type-specific gene expression analysis. This is, in part, because many genes are known to be part of this GO term. To study biological processes in which fewer genes are involved, we subsequently studied enriched GO terms with the highest odds ratios, i.e. with the largest difference in expected versus actual count. In this analysis, the endodermis is the only cell type that has a GO term that is enriched based on the up-regulation of more than five genes: suberin biosynthesis (GO:0010345) (Table 2). Suberin is a hydrophobic polymer deposited between the primary cell wall and the plasma membrane of endodermal cells. There suberin, together with the Casparian strip, block free movement of water and nutrients into the endodermis and consequently the innermost cell layers of the Arabidopsis root (Barberon, 2017; Geldner, 2013). Like the formation of lateral roots and root hairs, the production of suberin is affected by nutrient availability (Barberon et al., 2016). In addition, recent data suggest that suberin and the Casparian strip are involved in the interaction between plants and root-associated commensals, and soil-borne phytopathogens (Froschel et al., 2020; Salas-Gonzalez et al., 2021).

**Table 2.**
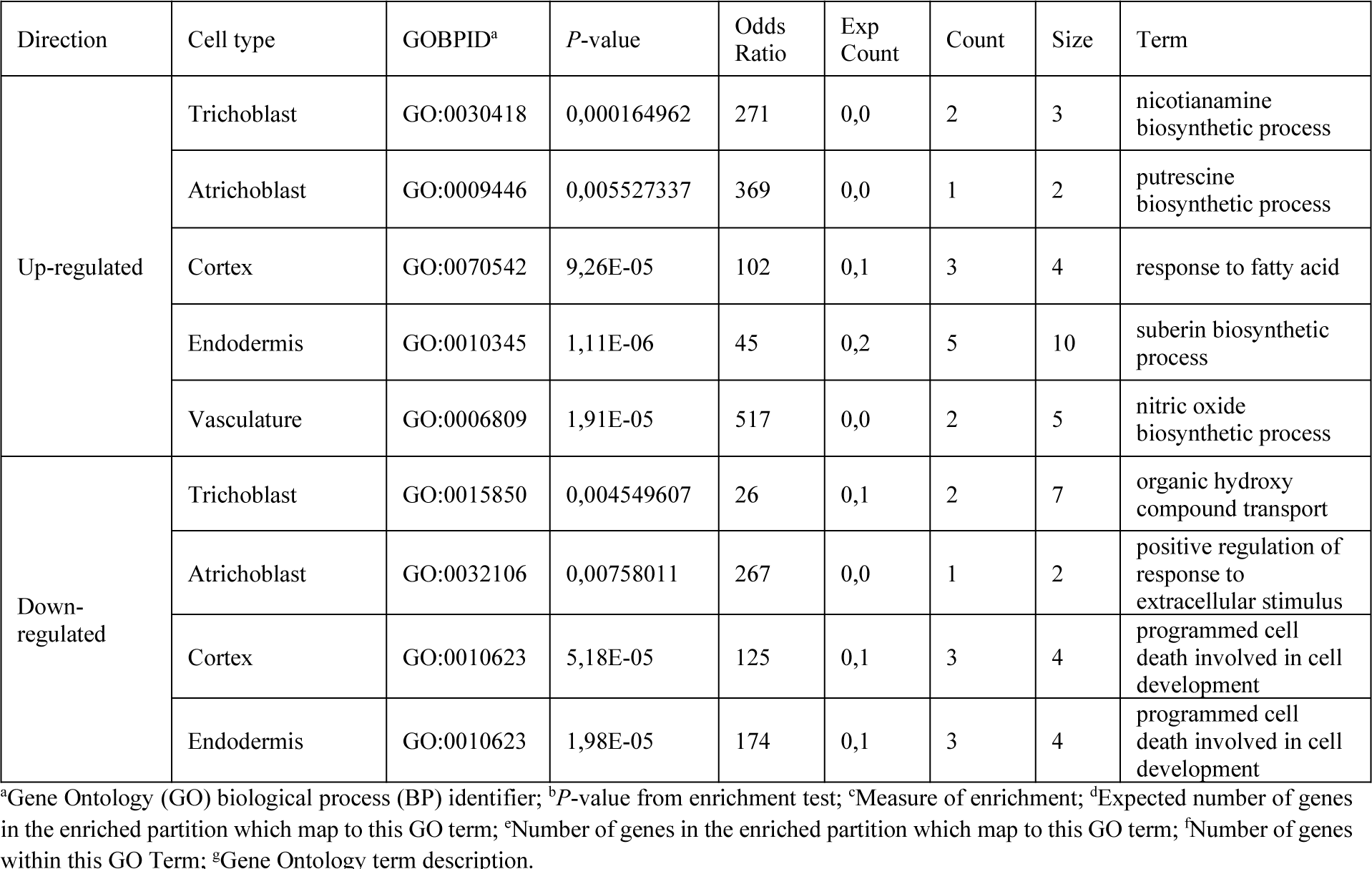
Top GO terms enriched in the WCS417-affected gene lists based on odds ratio.

We analyzed the effect of WCS417 on genes related with suberin production, such as MYB transcription factors suggested to activate suberin biosynthesis (Kosma et al., 2014; Lashbrooke et al., 2016; Shukla et al., 2021), and enzymes involved in suberin biosynthesis, including β-KETOACYL-CoA-SYNTHASEs (KCSs), fatty acid cytochrome P450 oxidases (CYP86A1 and CYP86B1), FATTY ACUL-CoA REDUCTASEs (FARs), GLYCEROL-3-PHOSPHATE SN2-ACYLTRANSFERASEs (GPATs) as well as transporters such as the ATP-binding cassette (ABC) transporter proteins (Barberon, 2017; Panikashvili et al., 2010; Vishwanath et al., 2015; Yadav et al., 2014). As expected, based on the available literature and our GO term analyses, spatial gene expression patterns show that suberin biosynthesis is primarily restricted to the endodermis and is significantly induced by WCS417 (Figure 6A-B). To validate the induction of suberin biosynthesis we imaged the transgenic plant line *GPAT5_pro_::mCITRINE-SYP122,* a reporter for suberin deposition, and stained roots with fluorol yellow to visualize suberin (Barberon et al., 2016). Consistent with our transcriptomic data, the *GPAT5* promoter is active specifically in the endodermis and is stimulated upon root colonization by WCS417 (Figure 6C). Additionally, WCS417 colonization led to an increase of suberin in the endodermis as quantified by a decreased distance from the root tip to the continuously suberized root zone (Figure 6D-E).

**Figure 6.**
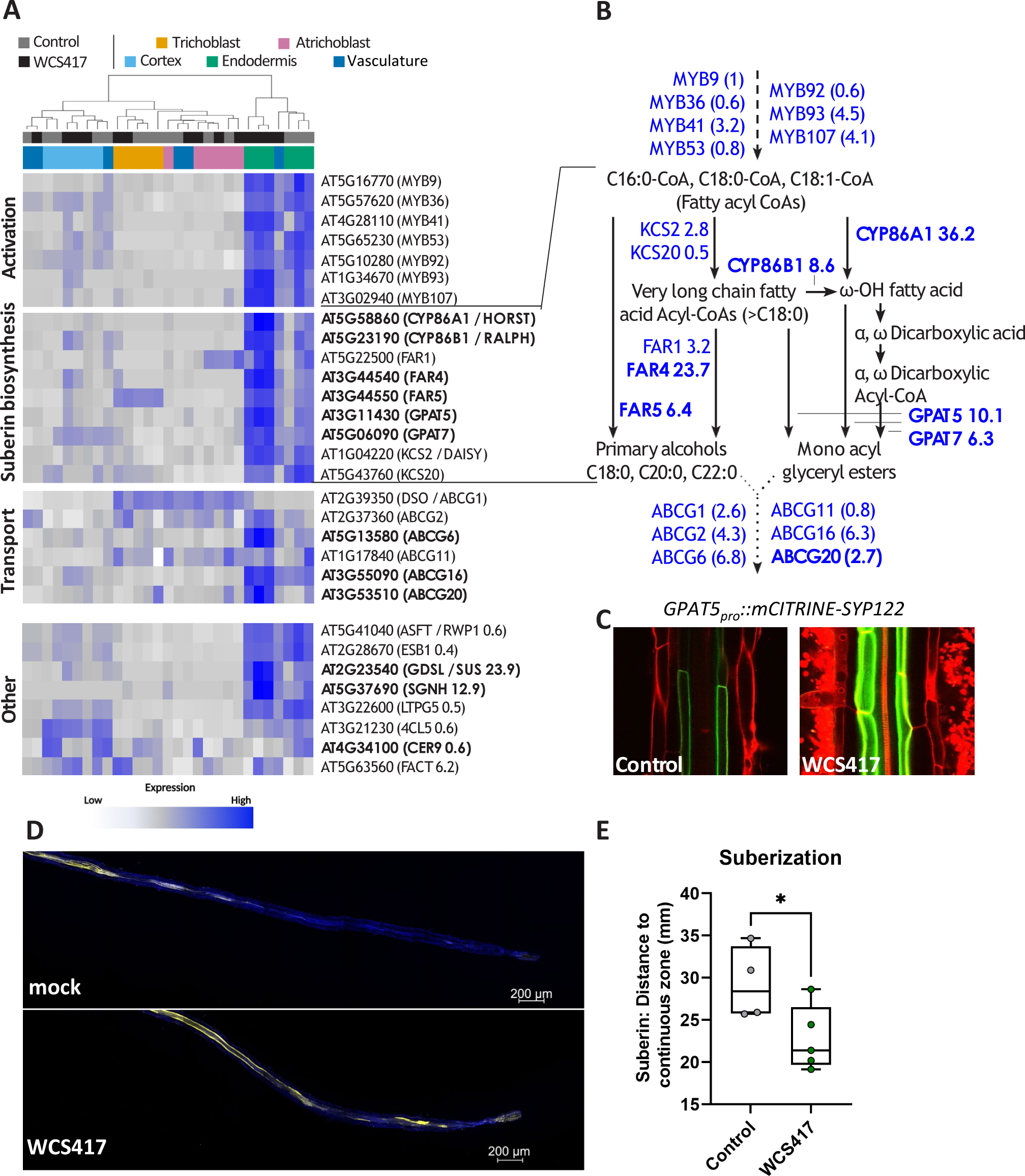
WCS417 induces suberization of the endodermis. **A**) Heatmap of the expression of genes known to be involved in suberin biosynthesis (GO:0010345, suberin biosynthetic process; Lashbrooke, 2016; Vishwanath, 2015). Heatmap is scaled per row (gene). Genes that are significantly up-regulated (logFC > 2, FDR < 0.1) by WCS417 in the endodermis are shown in bold. No significantly down-regulated genes were found in the endodermis. **B**) Overview of the suberin biosynthesis pathway, its activation and suberin monomer transport out of the cell, adapted from (Vishwanath *et al*. 2015). Genes known to be involved in these processes are shown in blue, fold changes as found in our dataset in response to WCS417 are shown. Statistical significantly differentially expressed genes (FDR < 0.1) are depicted in bold. Dashed lines show activation, solid lines show compound conversions, dotted lines show transport processes. **C**) Expression pattern of *GPAT5pro::mCITRINE-SYP122* in the Arabidopsis root, 2 days after inoculation with 105 CFU · ml-1 WCS417 or without treatment (control). Representative confocal images are shown. **D-E**) Suberization in roots of 7-d-old Arabidopsis at 2 d after they were transferred in Hoagland plates containing WCS417. Suberin was visualized using fluorol yellow staining and quantified as the distance from the root tip to the continuous zone of suberization in roots of Arabidopsis (n = 4-5). Representative confocal images are shown.

### Root endodermal barriers have a role in colonization by WCS417 and the subsequent activation of defense responses

Based on the increased suberization following colonization by WCS417 (Figure 6), we hypothesized that suberin and the Casparian strip might play a role in the interaction between Arabidopsis and WCS417. To test this, we grew wild-type plants and *myb36-2/sgn3-3* mutants, with developmentally delayed and reduced endodermal barriers (Salas-Gonzalez et al., 2021) and performed experiments with WCS417 similar to those described above for root hair mutants. The effects of WCS417 on wild-type and endodermal barrier mutant plants were similar in terms of shoot growth promotion (Figure 7A and Figure S3A). This was not the case for primary root length and lateral root formation, with WCS417 having a more pronounced effect on *myb36-2/sgn3-3* plants as compared to wild-type plants (Figure 7B-C and Figure S3B-C). Next, we tested the colonization levels of WCS417 on roots of wild-type and mutant plants.

**Figure 7.**
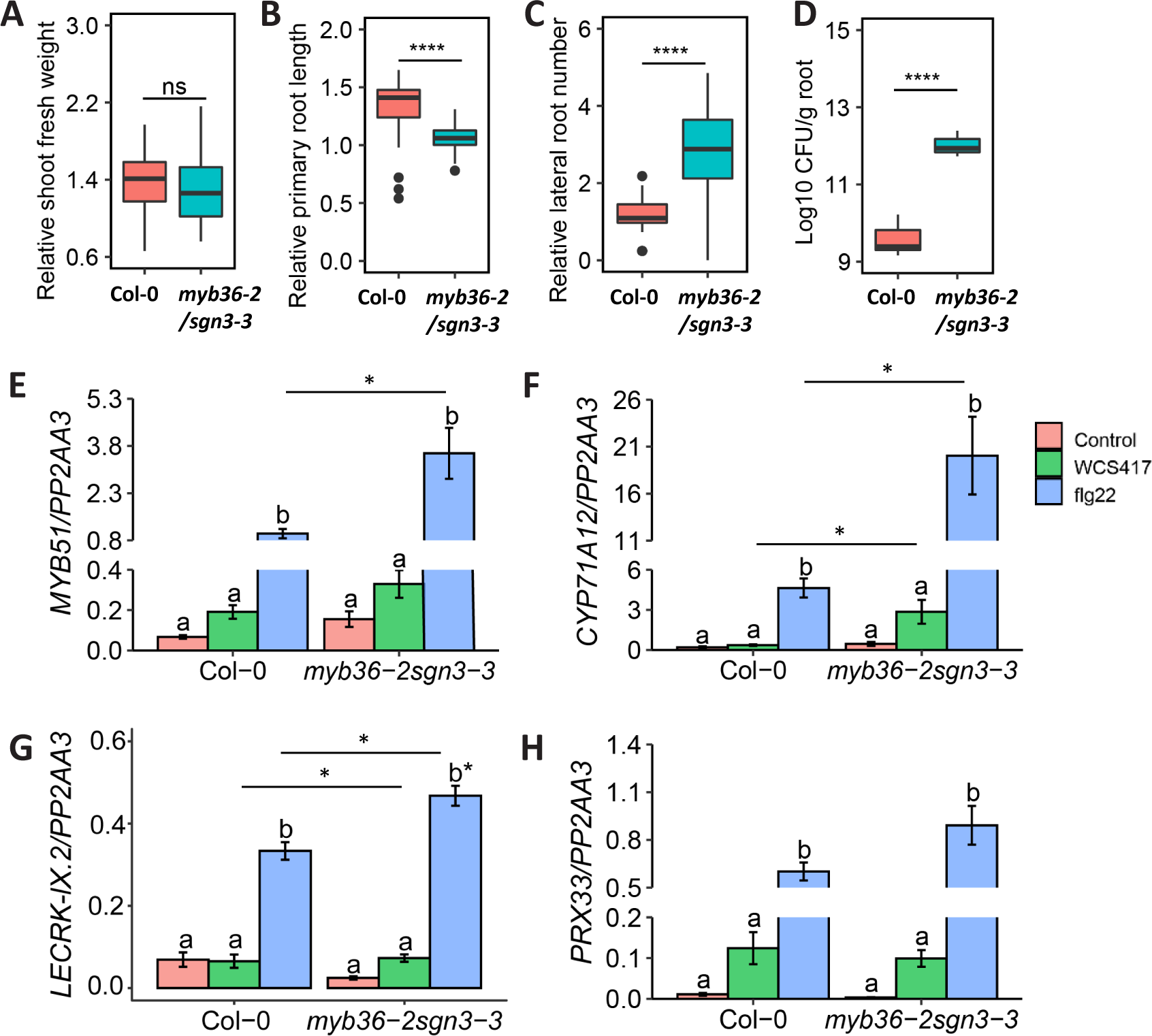
Root endodermal barrier integrity is needed for balanced interaction with WCS417. Relative **A**) shoot fresh weight, **B**) primary root length, **C**) lateral root number, and **D**) colonization levels of WCS417 on roots of Col-0 and *myb36- 2/sgn3-3* at 7 days after seedlings were transferred to plates with Hoagland medium (0% sucrose) containing 105 CFU · ml-1 WCS417. Different letters indicate statistically significant differences across genotypes (One-way ANOVA, Tukey’s test; *P* < 0.05). In the case of growth parameters n= 30 and in case of root colonization n= 6. (**E – H**) Expression levels of *MYB51*, *CYP71A12*, *LECRK-IX.2* and *PRX33* as quantified by qRT-PCR. Expression was tested in roots of 8-day-old seedlings at 6 h after inoculation with WCS417 (OD600= 0.1, 108 CFU · ml-1) or treated with 1 μM flg22. Error bars represent SEM. Different letters represent statistically significant differences among Control, WCS417 and flg22 in same plant genotype (One-way ANOVA, Tukey’s test; *P* < 0.05, n= 3-4). Asterisks indicate significant differences across genotypes that received the same treatment (WCS417 or flg22) (Student’s *t*-test; * *P* < 0.05, \*\**P* < 0.01, \*\*\**P* < 0.001,\*\*\*\**P* < 0.0001; ns, not significant).

Remarkably, WCS417 colonization levels were much higher (almost 100 times) on *myb36- 2/sgn3-3* roots as compared to the wild-type (Figure 7D) indicating that disruption of endodermal barriers greatly affects the interaction with WCS417. To further corroborate this observation, we studied the expression of MTI markers *MYB51, CYP71A12*, *PRX33* and *LECRK-IX.2* in roots of wild-type and *myb36-2/sgn3-3* treated with WCS417 and flg22 (Figure 7E-H). For all genes tested, there was a stronger response to flg22 in the endodermal barrier mutant, indicating that increased permeability of the endodermis makes roots more responsive to MAMPs. This is consistent with recent findings showing that flg22 could reach the inner cell types of plants with dysfunctional endodermal barriers and activate stonger expression of MTI markers (Zhou et al., 2020). The roots of the endodermal barrier mutant also produced a stronger induction of *CYP71A12* following treatment with WCS417 (Figure 7F), suggesting that the increased colonization of these roots (Figure 7D) can lead to stronger root defense responses, probably via diffusion of WCS417 MAMPs into deeper root layers.

## Discussion

### Creating a spatial map of gene expression changes in response to WCS417

The ‘hidden half’ of plants, the root system, is of crucial importance when breeding for plants that are drought tolerant or better able to grow under nutrient-limiting conditions (Rogers and Benfey, 2015; Koevoets *et al*., 2016). Additionally, the root surface lies at the interface between plants and beneficial soil micro-organisms, which increase plant growth and health (Lugtenberg and Kamilova, 2009; Pieterse *et al*., 2014; Bakker *et al*., 2018). To better understand the response to beneficial bacteria at the level of individual cell types, we studied gene expression changes in five Arabidopsis root cell types after colonization with WCS417 (Figures 2-3). The total number of DEGs identified across the five cell types is approximately ten-fold greater than the number identified in the sorted whole root control. A similar increase in detection power of cell-type-specific versus whole root transcriptomic analyses was obtained previously when examining the Arabidopsis root response to salt, iron deficiency, and nitrogen (Dinneny *et al*., 2008; Gifford *et al*., 2008).

We show that the increased sensitivity can be traced to the cell-type-specific nature of the root response to WCS417. The five cell types differ in their response to WCS417 both quantitatively, with large differences in the number of DEGs, and qualitatively, with little overlap in DEGs between cell types (Figure 3). This supports previous studies on cell-type-specific gene expression changes in response to both abiotic and biotic stresses and refutes the concept of a global stress response (Dinneny et al., 2008; Gifford et al., 2008; Iyer-Pascuzzi et al., 2011; Rich-Griffin et al., 2020a; Walker et al., 2017). The little overlap in DEGs between cell types results in many genes that are up- or down-regulated in only a single cell type, and the majority of these genes are not identified as differentially expressed in our whole-root controls (Tables 1 and S3-S4). Thus, cell-type-specific transcriptional profiling is more sensitive than whole-tissue transcriptional profiling because it detects cell-type-specific DEGs that are otherwise hidden.

### Cell-type specific signatures of the WCS417-Arabidopsis interaction

The number and type of DEGs in our spatial map uncovered two interesting patterns: 1) the cortex and endodermis respond most strongly to WCS417 in terms of the number of DEGs, and 2) the number, type and basal expression level of DEGs in the two epidermal cell types is distinct (Figure 4). The strong response of the cortex and endodermis is surprising, as these cell types are likely not in direct contact with WCS417. Previous studies, however, demonstrated the ability of MAMPs to reach the cortex and the endodermis (Zhou et al., 2020), and mount MTI responses (Wyrsch et al., 2015). Also timing likely plays a role, as cell-type-specific transcriptional profiling of the Arabidopsis root response to flg22 showed that the epidermis responded as strongly as the cortex at two hours post inoculation (Rich-Griffin et al., 2020a). Possibly, the epidermis responds strongly at first and down-regulates its response by two days after inoculation, while the cortex and endodermis maintain or increase their response over that time frame, to restrict continuous and unwanted activation of the outer cell types exposed to the microbe-rich environment. We observed enrichment of processes related to decreased cell wall biogenesis in these inner cell types specifically. This might allow these cells to lose volume which is required for lateral root initiation and outgrowth (Vermeer *et al*., 2014; Stoeckle *et al*., 2018), and might explain the observed increase in lateral root formation in WCS417-exposed roots (Stringlis et al., 2018a; Zamioudis et al., 2013). A time-series experiment could further elucidate the timing and magnitude of these spatially-separated responses, while future studies on the responsiveness of younger and older parts of the root to different stimuli could provide further evidence on how roots contribute to plant homeostasis in microbe-rich and stress- abundant environments.

In addition to decreased cell wall biogenesis, we show increased expression of genes involved in suberin biosynthesis in the endodermis (Figure 6). We confirmed increased suberization of the endodermis by visualization of *GPAT5_pro_*::*mCITRINE-SYP122* activity and suberin staining (Figure 6). Suberin and the Casparian strip are essential for protecting the inner root tissues from the surrounding soil environment (Barberon, 2017). Suberin displays plasticity to nutrient stresses such as iron deficiency (Barberon et al., 2016), but it can also be modulated in response to beneficial and pathogenic members of the microbiome (Froschel et al., 2020; Kashyap et al., 2022; Salas-Gonzalez et al., 2021). It is probable that both beneficial and pathogenic microorganisms manipulate the functioning and deposition of endodermal barriers to achieve sufficient colonization of the root and access to root-derived sugars. Indeed, previous research has shown that Arabidopsis activates the iron deficiency response upon root colonization by WCS417 (Verhagen et al., 2004; Zamioudis et al., 2014; Zamioudis et al., 2015). This response is normally activated when plants experience a shortage of iron and results in a decreased deposition of suberin to facilitate iron uptake (Barberon et al., 2016), suggesting that WCS417 modulates nutrient availability or use within the plant. Our data further suggest that this might be a transient response during colonization, since at 48 h after colonization we observed increased, rather than decreased, suberization in the roots, suggesting that plants adapt to the interaction with WCS417 and re-seal the endodermis to avoid unwanted effects. This hypothesis is supported by our experiment with the *myb36-2*/*sgn3-3* double mutant with dysfunctional endodermal barriers. This mutant is colonized to a higher degree by WCS417 and has elevated expression of the MTI marker gene *CYP71A12* compared to wild-type plants (Figure 7). Additionally, in this mutant three of four MTI marker genes show increased expression in response to flg22, further confirming the role of this barrier in MTI sensitivity and in fine-tuning growth and defense. Therefore, for an optimal interaction with WCS417, Arabidopsis needs a functional endodermal barrier to prevent limitless bacterial proliferation on the root.

Another interesting observation was the distinct transcriptomic behavior of the two epidermal cell types under untreated conditions (Figure 4). These differences turned out to be at least in part due to functional specialization of trichoblasts in nutrient uptake and atrichoblasts in basal defense gene activation. Such functional specialization might benefit plants to deal efficiently with both biotic and abiotic stresses simultaneously. This is consistent with previous findings showing increased expression of growth-related genes in progressively more differentiated trichoblasts (Denyer *et al*., 2019) and specialization of trichoblasts in nutrient uptake (Vert *et al*., 2002; Tanaka *et al*., 2014; Zamioudis *et al*., 2014). Based on our findings, it is tempting to speculate that root hairs could act as antennae perceiving environmental signals and informing plants to adapt their growth and development to an upcoming interaction. Literature supports this, since in other plant species root hairs are colonization hotspots for Rhizobia (Poole et al., 2018), the formation and pattern of root hairs are responsive to nutrient stresses (Vissenberg et al., 2020), root hairs mediate exudation of iron-mobilizing coumarins (Robe et al., 2021), and barley mutant plants with contrasting root hair characteristics accommodate distinct root-associated microbial communities (Robertson-Albertyn et al., 2017).

### Concluding remarks

We created a spatial map of gene expression changes induced in the Arabidopsis root in response to colonization by the beneficial bacterium WCS417. Our dataset uncovers localized, cell-type-specific gene expression patterns that otherwise remain hidden in global analyses of gene expression and that correspond to observed root architectural changes. We demonstrate a role for root hairs and endodermal barriers in the interaction between roots, WCS417 and microbial MAMPs. In addition, further mining of our dataset will enable other researchers to determine the spatial pattern of microbe-induced expression of genes of interest.

## Methods

### Plant material and growth conditions

#### FACS experiment

Arabidopsis accession Columbia-0 (Col-0) and transgenic Col-0 with the *COBRA- LIKE9_pro_:GFP* (Brady et al., 2007a; Brady et al., 2007b), *WEREWOLF_pro_:GFP* (Lee and Schiefelbein, 1999), *315_pro_:GFP* (Lee et al., 2006), *SCARECROW_pro_:GFP* (Wysocka-Diller et al., 2000), or *WOODENLEG_truncated_pro_:GFP* construct (Mahonen et al., 2000) were grown as described previously (Dinneny et al., 2008). Briefly, seeds were liquid sterilized in 50% bleach and stratified by incubation at 4°C for 2 d. Sterilized seeds were plated in two dense lines of three seeds thick each on nylon mesh (Nitex Cat 03-100/44, Sefar) on sterile 1 × MS (Murashige and Skoog (1962)) 1% sucrose plates. Plates were sealed with Parafilm and placed vertically in long day conditions (22°C; 16 h light, 8 h dark) for a total of 7 d.

#### Microscopy for suberin localization

Col-0 seeds were surface sterilized (Van Wees et al., 2013) and sown on plates containing agar-solidified Hoagland medium with 1% sucrose and pH was adjusted to 5.5 (Stringlis et al., 2018a). After 2 d of stratification at 4°C, the plates were positioned vertically and transferred to a growth chamber (22°C; 10 h light, 14 h dark; light intensity 100 μmol · m^−2^ · s^−1^). When 5-d-old, seedlings were transferred to agar-solidified Hoagland plates without sucrose (0.75% agar) where *Pseudomonas simiae* WCS417 (WCS417) was mixed in the medium based on the protocol developed by Paredes et al. (2018). After 2 d of Arabidopsis-WCS417 interaction, Fluorol yellow (FY) staining of roots was performed as described before (Kajala et al., 2021; Lux et al., 2005).

#### Colonization and growth promotion experiments with endodermal barrier and root hair mutants

Col-0 seeds and mutants in Col-0 background: *myb36-2/sgn3-3* (Reyt et al., 2021), *cpc-1* and *ttg1* (Wada et al., 1997; Walker et al., 1999) were surface sterilized and sown on agar-solidified Hoagland plates (as before). When 7-d-old, seedlings were transferred to agar-solidified Hoagland plates with 0% sucrose where WCS417 was mixed in the medium (as before). Seven days later the shoots of the seedlings were weighed using an analytical scale, photos were taken to analyze root growth and development (via Image J) and colonization of WCS417 on roots was assessed (Paredes et al., 2018).

#### Analysis of gene expression in endodermal barrier and root hair mutants

For testing root transcriptional responses to WCS417 and flg22, plants were grown and treated based on a protocol developed by Stringlis et al. (2018a). Briefly, uniform 9-day-old seedlings were transferred from MS agar plates to six-well plates (ø 35 mm per well) containing liquid 1 × MS with 0.5% sucrose, after which they were cultured for 7 more days under the same growth conditions. One day before treatment with either WCS417 or flg22, the medium of each well was replaced with fresh 1 × MS medium with 0.5% sucrose. At 6 h after treatment with WCS417 or 1 μM flg22 (GenScript), roots were flash frozen in liquid nitrogen for downstream gene expression analysis.

### WCS417 treatment

#### FACS experiment

Plants were inoculated with bacteria 5 d after being placed in long-day conditions using a slightly adapted version of a previously published protocol (Zamioudis *et al*., 2015). Briefly, rifampicin-resistant WCS417 was streaked from a frozen glycerol stock onto solid King’s medium B (KB) (King *et al*., 1954) containing 50 µg · ml^-1^ rifampicin and grown at 30°C overnight. One day before plant treatment, a single colony from the plate was put in liquid KB with rifampicin and grown in a shaking incubator at 30°C overnight. The following morning, the suspension reached an OD_600_ value between 0.6 and 1.0 (OD_600_ of 1.00 is equal to 10^9^ colony-forming units (CFU) · ml^-1^), after which the bacteria were washed twice with 10 mM MgCl_2_.

To decide which bacterial concentration to add to the plants, the washed bacteria were resuspended in 10 mM MgCl_2_ to a final density ranging from 10^1^ to 10^8^ CFU · µl^-1^. Two horizontal lines of either 10 µl of 10 mM MgCl_2_ or 10 µl of one of the bacterial suspensions were applied per 1 × MS 1% sucrose plate. Five-day-old Col-0 seedlings were transferred on their mesh onto these plates. Seedlings were transferred such that the roots of the seedlings were on top of the bacteria. Finally, all plates were resealed with Parafilm and left to grow in long-day conditions. At 2 and 7 d after treatment ten plants from each treatment were randomly picked and removed from the plate. The total number of emerged lateral roots was counted under a stereo microscope. ImageJ was used to determine primary root length per plant from images made with a scanner.

Based on the results of this trial, we chose a density of 10^6^ CFU · µl^-1^, amounting to 10^7^ CFU per row of plants, for the sorting experiment (see below). Wild-type Col-0 plants and plants of each of the five transgenic lines were exposed to this bacterial density after 5 d of plant growth as described above and incubated in long-day growth conditions for an additional 2 d.

#### Suberin staining experiment and colonization, assessment of growth and gene expression of endodermal and root hair mutants

For the rest of experiments, WCS417 was prepared and applied based on previously established protocols (Paredes et al., 2018; Stringlis et al., 2018a). WCS417 was cultured at 28°C on KB agar plates supplemented with 50 μg · ml^-1^ of rifampicin. After 24 h of growth, cells were collected in 10 mM MgSO_4_, washed twice with 10 mM MgSO_4_ by centrifugation for 5 min at 5000 *g*, and finally resuspended in 10 mM MgSO_4_. For suberin staining and growth/colonization experiments of Col-0 and mutant seedlings, WCS417 was mixed in Hoagland agar plates without sucrose in a concentration of 10^5^ CFU · ml^-1^. This mix was then poured in plates and the seedlings were transferred in the plate once solidified.

For qRT-PCR gene expression analysis of Col-0 and respective mutants, WCS417 bacteria were added in each well to a final OD of 0.1 at 600 nm (10^8^ CFU · ml^-1^).

### Fluorescence-activated cell sorting (FACS)

After a total of 7 d of growth, roots were cut from the shoot with a carbon steel surgical blade. Whole roots of Col-0 destined for the unsorted control were immediately frozen in liquid nitrogen in an Eppendorf tube. For the other samples, all to be put through a cell sorter, roots were cut twice more and the root pieces from 4 - 6 plates were collected and protoplasted as described previously (Birnbaum *et al*., 2003, 2005). Briefly, they were placed in a 70-µm cell strainer submerged in enzyme solution (600 mM mannitol, 2 mM MgCl_2_, 0.1% BSA, 2 mM CaCl_2_, 2 mM MES, 10 mM KCl, pH 5.5 with 0.75 g cellulysin and 0.05 g pectolyase per 50 ml). Roots were mixed in the strainer at room temperature (RT) on an orbital shaker to dissociate protoplasts. After one hour, the suspension surrounding the strainer, containing the protoplasts, plus a few roots to pull the protoplasts down, were pipetted into a 15-ml conical tube and spun at 200 *g* for 6 min at RT. The top of the supernatant was pipetted off and the remaining solution resuspended in 700 µl of the protoplasting solution without enzymes (600 mM mannitol, 2 mM MgCl_2_, 0.1% BSA, 2 mM CaCl_2_, 2 mM MES, 10 mM KCl, pH 5.5). This suspension was filtered successively through a 70-µm cell strainer and a 40-µm strainer. The filtrate was finally collected in a cell sorting tube and taken to the cell sorter (Astrios, Beckman Coulter) at RT.

Protoplasts sorted by the machine were collected into RLT buffer (Qiagen) with β- mercaptoethanol. The samples were immediately placed on dry ice to inhibit RNA degradation. Samples were stored at -80°C until RNA isolation.

### RNA isolation and sequencing

#### FACS experiment

Whole root tissue for the unsorted control was lysed by grinding with a liquid-nitrogen-cooled mortar and pestle. RNA was isolated with the RNeasy Plant Mini Kit (Qiagen) for the 6 unsorted whole root samples and for 6 out of 8 sorted whole root samples. RNA from the remaining two sorted whole root samples and all cell type-enriched samples were isolated with the Micro Kit (Qiagen). RNA concentration was checked with a Qubit Fluorometer (Thermo Scientific) and RNA integrity was assessed with a Bioanalyzer (Agilent Technologies). Subsequently, RNA libraries were made from samples with RNA integrity number (RIN) values above six. All libraries were made with the NEBNext Ultra RNA Library Prep Kit for Illumina (NEB). RNA for the 6 control unsorted (whole root) and the first 6 control sorted samples were poly-A selected using Dynal Oligo-dT beads. These 12 libraries were generated using 100 ng of total RNA. The remaining libraries were generated from total RNA selected populations, 50 ng of total RNA was used as starting material for all sorted library preparations. Libraries were sequenced on an Illumina HiSeq 2500 using 50 base pair Single-Read (Duke University Sequencing Core). Three biological replicates were performed for each sample type and condition, except for the sorted control, for which we performed 4 biological replicates.

#### qRT-PCR experiment of Col-0 and mutants

For the qRT-PCR experiment, roots of Col-0 and mutants were collected in 4 replicates at 6 h after treatment with live WCS417 cells or flg22. Roots of untreated Col-0 and mutant seedlings were collected at the same time point (as controls). Each of the 4 biological replicates per treatment consisted of 10-12 pooled root systems. After harvest, root samples were snap-frozen in liquid nitrogen and stored at −80°C. Arabidopsis roots were homogenized using a mixer mill (Retsch) set to 30 Hz for 45 s. RNA extraction was performed with the RNeasy Plant Mini Kit (Qiagen). RNA concentration was checked with a Qubit Fluorometer (Thermo Scientific). For qRT-PCR analysis, DNase treatment, cDNA synthesis and PCR reactions and subsequent analysis were performed as described by Stringlis et al. (2018b). Primer sequences for the reference gene *PP2AA3* and the MTI marker genes *MYB51*, *CYP71A12, LECRK-IX.2 and PRX33* are listed in Table S1.

### Data analysis

The reads generated by Illumina sequencing were pseudoaligned to the TAIR10 cDNA database (Lamesch *et al*., 2012) using Kallisto (v0.43.0) with 100 bootstraps and default settings (Bray *et al*., 2016). The percentage of aligned reads is lower for the 12 samples that were poly-A selected using Dynal Oligo-dT beads because of a high number of rRNA sequences. This is probably due to differences in the bead-selection procedure and greater amount of RNA used as starting material. We do not expect this to interfere with our analyses, as the number of expressed genes in these samples is in the same range as previously published data in this species. The resulting transcript counts were subsequently summarized to the gene level with tximport (v1.2.0) (Soneson *et al*., 2015). One bacteria-exposed sample enriched for trichoblasts was excluded from further analyses because of low coverage. Only genes with a count per million (cpm) greater than two in all samples were kept for the remaining analysis. The counts per gene of the remaining samples and genes were used to generate a digital gene expression list (DGE list) in EdgeR (v3.16.5) (Robinson *et al*., 2010). A generalized linear model (glm) was fit using a negative binomial model and quasi-likelihood (QL) dispersion estimated from the deviance with the glmQLFit function in EdgeR. DEGs were then determined by comparing the bacteria-exposed and the non-exposed samples with the glmQLFTest (FDR < 0.1; -2 < log_2_FC > 2). GO term analysis was performed in R based on the genome wide annotation for Arabidopsis within org.At.tair.db (Carlson M, 2018) with the program GOstats (Falcon and Gentleman, 2007).

### Fluorescence microscopy

Approximately 20 sterilized and vernalized seeds of the *COBL9_pro_:GFP*, *WER_pro_:GFP*, *315_pro_:GFP*, *SCR_pro_:GFP*, and *WOL_truncated_pro_:GFP* transgenic lines were sown on a 1 × MS 1% sucrose plate and placed in long-day conditions. After 5 d either 10^5^ WCS417 cells in 10 µl MgCl_2_ or sterile 10 µl MgCl_2_ was added to each root. GFP localization was observed once per day in 5, 6 and 7-day-old seedlings with a 510 upright confocal microscope with a 20x objective (Zeiss).

*GLYCEROL-3-PHOSPHATE ACETYLTRANSFERASE 5_pro_:mCITRINE-SYP122* (*GPAT5_pro_::mCITRINE-SYP122*) plants (Barberon et al., 2016) were grown on MS plates in long-day conditions. After 5 d, 10^5^ bacteria 10 µl MgCl_2_ were inoculated onto each root tip. Fluorescence was checked with a 510 upright confocal microscope (Zeiss) at 2 d after inoculation.

For FY staining of suberin, Col-0 seeds were sown on Hoagland plates 1% sucrose and placed in short-day conditions. When 5-days-old, seedlings were transferred in Hoagland medium without sucrose mixed with 10^5^ CFU · ml^-1^ WCS417. After 2 d, roots were washed in MQ, separated from the leaves and 5 roots were added in each well of 6-well plates and incubated in FY088 (0.01% w/v, dissolved in lactic acid) for 1 h at RT in darkness, rinsed three times with water (5 mins per wash), and counterstained with aniline blue (0.5% w/v, dissolved in water) for 1 h at RT in darkness. Roots were mounted with 50% glycerol on glass slides and kept in dark until observation. Confocal Laser Scanning microscopy was performed on a Zeiss LSM 700 laser scanning confocal microscope with the 20X objective and GFP filter (488nm excitation, 500-550nm emission). To quantify the suberization pattern, the distance from the root tip to the start of continuous suberization was determined with ImageJ (v1.53g).

### Data availability

The raw RNA-Seq read data are deposited with links to BioProject accession number PRJNA836026 in the NCBI BioProject database.

## Funding

This research was funded in part by the Netherlands Organization of Scientific Research through ALW Topsector Grant no. 831.14.001 (E.H.V.), by a postdoctoral fellowship from the Jane Coffin Childs Memorial Fund for Medical Research (L.M.L.), by the NIH (5R01-GM- 043778), the NSF (MCB-06-18304), the Gordon and Betty Moore Foundation and the Howard Hughes Medical Institute (P.N.B.), by a postdoctoral fellowship from the Research Foundation Flanders (FWO 12B8116N) (R.d.J.), the China Scholarship Council (CSC) scholarship no. 201908320054 (JZ) and scholarship no 202006990074 (JY), the Technology Foundation Perspective Program “Back2Roots” Grant no. 14219 (C.M.J.P.), the ERC Advanced Grant no. 269072 of the European Research Council (C.M.J.P.), and the NWO Gravitation Grant no. 024.004.014 (I.A.S and C.M.J.).

## Author contributions

Conceptualization: E.H.V., L.M.L., P.N.B., C.M.J.P., I.A.S, and R.d.J., Methodology: E.H.V., L.M.L., Formal Analysis: E.H.V. and R.d.J., Investigation: E.H.V., L.M.L., J.Z., J.Y. and I.A.S., Writing – Original Draft: E.H.V, L.M.L. C.M.J.P., I.A.S. and R.d.J., Writing – Review & Editing: L.M.L. I.A.S., P.N.B., C.M.J.P. and R.d.J., Visualization: E.H.V., L.M.L, J.Z., J.Y., R.d.J. and I.A.S. Funding Acquisition: E.H.V., L.M.L. C.M.J.P, P.N.B. and R.d.J.

## Acknowledgements

The authors want to thank Niko Geldner for the *GPAT5_pro_*::*mCITRINE-SYP122* and *myb36- 2/sgn3-3* lines, Christian Dubos for *cpc* and *ttg1* lines, Cara M. Winter for help with *GPAT5_pro_::mCITRINE-SYP122* microscopy, and Rosa Toonen for the experiment involving suberin staining and confocal microscopy. There is no conflict of interest to declare.

---

**Figure S1.**
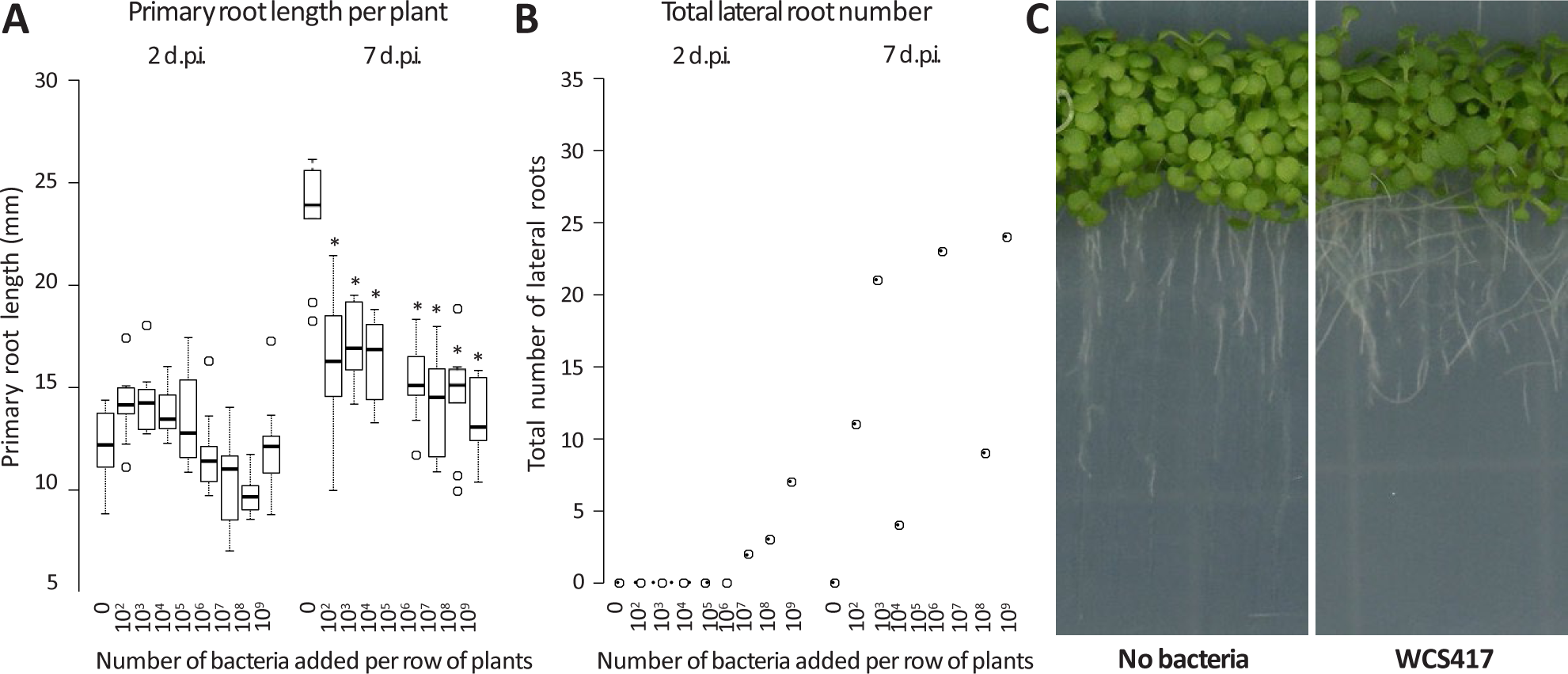
The effect of WCS417 on root system architecture of Arabidopsis. A) Quantification of primary root length two- and seven-days post inoculation (d.p.i) with increasing numbers of WCS417 bacterial cells. Asterisks represent a significant difference compared to the control from the same time point (ANOVA, post hoc Tukey, p-value < 0.05). B) The total number of lateral roots of ten seedlings two and seven days after inoculation with increasing numbers of bacterial cells. C) Pictures of plants in the densely sown set-up six days after application of a mock solution or a solution containing 10^7^ bacterial cells per row of plants.

**Figure S2.**
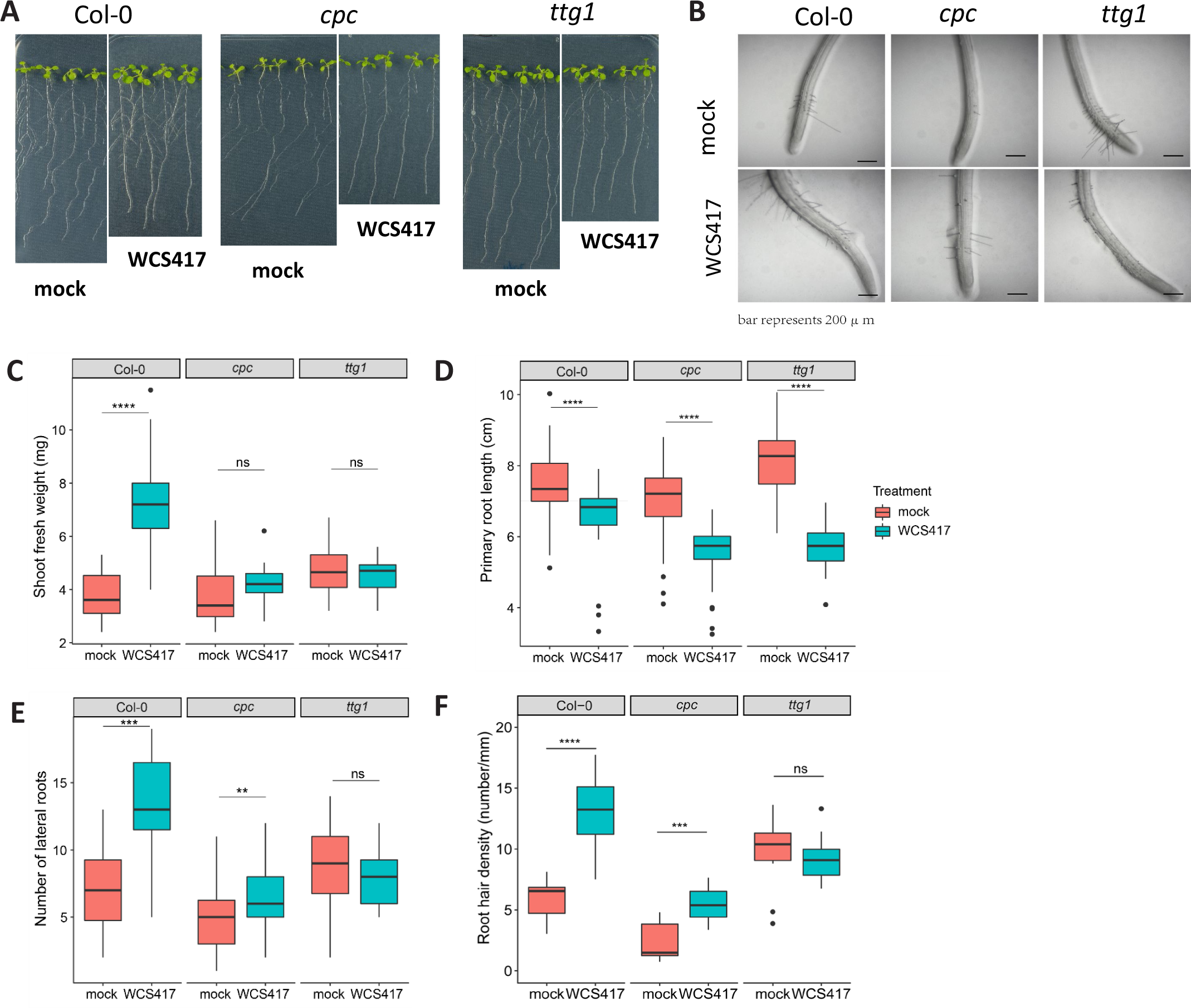
Growthpromotiondata of Col-0 and root hair mutants in response to WCS417. Representative **A**) macroscopic photos of Col-0, *cpc* and *ttg1* seedlings and **B**) microscopic photos of the area 1 cm above root tip of Col-0, *cpc* and *ttg1* seedlings at 7 days after seedlings were transferred to plates with Hoagland (0% sucrose) containing 10^5^ CFU of WCS417 per ml of medium. (**C – F**) Measurements of shoot fresh weight, primary root length, number of lateral roots and root hair density at 7 days after seedlings were transferred to plates with Hoagland (0% sucrose) containing 10^5^ CFU of WCS417 per ml of medium.. Asterisks indicate significant difference across genotypes with the same treatment (WCS417, flg22) (Student’s *t*-test; * *P* < 0.05, \*\**P* < 0.01, \*\*\**P* < 0.001,\*\*\*\**P* < 0.0001; ns, not significant). In the case of growth parameters n= 39– 40 and in case of root hair measurements n= 10.

**Figure S3.**
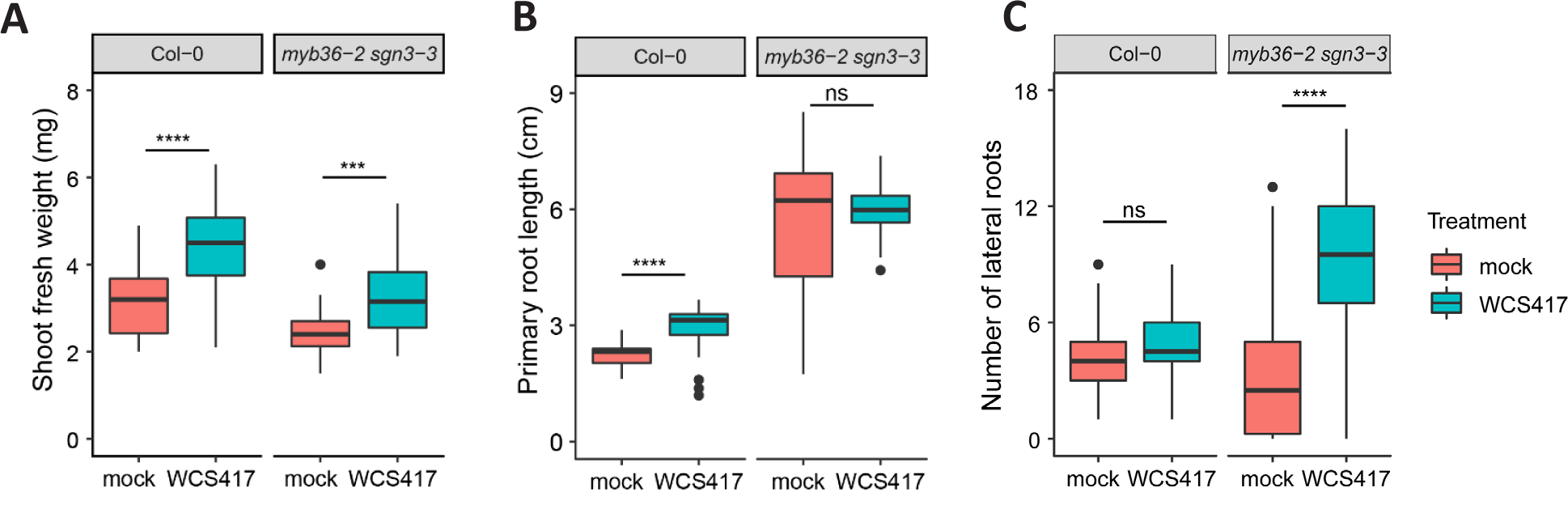
Growth promotion data of Col-0 and *myb36-2 sgn3-3* in response to WCS417. Measurements of **A**) shoot fresh weight, **B**) primary root length and **C**) number of lateral roots at 7 days after Col-0 and *myb36-2 sgn3-3* seedlings were transferred to plates with Hoagland (0% sucrose) containing 10^5^ CFU of WCS417 per ml of medium. Asterisks indicate significant difference across genotypes with the same treatment (WCS417, flg22) (Student’s *t*-test; \*\*\**P* < 0.001,\*\*\*\**P* < 0.0001; ns, not significant). n= 30

## Notes

### Competing Interest Statement

The authors have declared no competing interest.

